# Identification of transcriptome-wide cobalt chloride-induced hypoxia-responsive long noncoding RNAs regulated by cytoplasmic mRNA capping enzyme

**DOI:** 10.1101/2024.11.21.624636

**Authors:** Pinaki Ghosh, Safirul Islam, Chandrama Mukherjee

**Affiliations:** RNA Bio Lab, Institute of Health Sciences, Presidency University, Plot No. DG/02/02, Premises No. 14-0358, Action Area 1D, Kolkata 700156, India

**Keywords:** Long noncoding RNAs, CoCl_2_-induced hypoxia, mRNA capping enzyme, Cytoplasmic capping, uncapped RNA

## Abstract

Long non-coding RNAs (lncRNAs) are key regulators of gene expression in human cancers, influencing tumor microenvironment (TME). Hypoxia, a hallmark of solid tumors, elevates hypoxia-inducible factor 1α (HIF-1α), which regulates hypoxia-responsive lncRNAs (HRLs). These HRLs are essential in regulating gene expression under hypoxic conditions at both transcriptional and post-transcriptional levels. However, the mechanisms governing the post-transcriptional regulation of lncRNAs are not fully understood. It has been shown that the cytoplasmic pool of mRNA capping enzyme (cCE) can post-transcriptionally regulate its substrate uncapped mRNAs and lncRNAs in the cytoplasm, thereby preventing their degradation. Our previous study showed elevated expression of cCE under hypoxic conditions. This study aimed to determine whether cCE influences the transcriptome-wide HRLs in CoCl_2_-induced hypoxia in osteosarcoma U2OS cells. We identified 306 known lncRNAs with significant differential expression under hypoxic conditions, including 137 upregulated and 169 downregulated by RNA sequencing. Gene Ontology enrichment analysis revealed their distinct association with various cancers. Our investigation aimed to ascertain whether the cCE post-transcriptionally regulates these HRLs. Overexpression of a dominant negative form of cCE suggested cCE might post-transcriptionally regulate these HRLs. Future research should focus on understanding how cCE influences substrate HRLs, which play a crucial role in the bidirectional signaling circuit between TME and cancer cells under hypoxic stress as well as establishing cCE as a key factor that may alter cellular responses to hypoxia by stabilizing cCE-targeted lncRNAs.

## Introduction

The tumor microenvironment (TME) plays a crucial role in driving cancer progression by promoting cell growth and survival and contributing to the changes in disease characteristics [1]. Unrestrained tumor proliferation often leads to insufficient oxygen supply, resulting in hypoxia. Hypoxia, a defining characteristic of the TME, is prevalent in most solid tumors and is caused by imbalance in oxygen supply[2]. Hypoxic regions are a significant independent prognostic factor for human cancer, leading to more aggressive and therapeutically resistant tumor phenotypes[3]. The resulting changes in gene expression and subsequent proteomic alterations have profound effects on cellular and physiological functions, ultimately impacting patient prognosis[4]. Additionally, hypoxia contributes to the plasticity and heterogeneity of tumors, promoting a more aggressive and metastatic phenotype [5], [6], [7], [8]. The increased expression of hypoxia-inducible factor 1α (HIF-1α) plays a pivotal role in this process. HIF is a DNA-binding complex comprising two basic helix-loop-helix proteins from the PAS family: the constant HIF-1 β and either HIF-1 α or HIF-2 α[9]. In low oxygen conditions, the α/β heterodimer binds to a core pentanucleotide sequence (RCGTG) within the hypoxia response elements (HREs) of target genes[10], [11]. This interaction triggers an adaptive response to hypoxia, influencing tumor progression and metastasis[12].

The human genome is largely transcribed into non-coding RNAs (ncRNAs) that do not encode proteins. These ncRNAs are known to play critical roles in regulating the onset and progression of various types of cancer. A growing number of long non-coding RNAs (lncRNAs) have been identified to play a role in cellular stress responses, including those triggered by hypoxia, genotoxic stress, and oxidative stress[13]. Recent studies have underscored the significance of the non-coding genome in hypoxic tumor regions. Specifically, hypoxia-responsive long non-coding RNAs (HRLs) have been found to play key roles in regulating gene expression in hypoxic conditions at transcriptional, and post-transcriptional levels[14], [15]. However, the post-transcriptional control and stability of large lncRNAs is still not fully understood. Furthermore, lncRNAs are crucial in regulating epigenetic mechanisms within the chromatin structure, playing a significant role in maintaining chromatin architecture[16]. As these lncRNAs are crucial for gene regulation, gaining insight into their post-transcriptional regulation is essential for uncovering their functional significance.

5’ capping is a crucial post-transcriptional event that plays a significant role in determining the stability of both mRNAs and lncRNAs. The process of nuclear capping occurs co-transcriptionally, while cytoplasmic capping takes place post-transcriptionally. The eukaryotic RNAs transcribed by RNA polymerase II need a 5’ cap structure. This cap consists of an N7-methylated guanosine residue that is attached to the first nucleotide of the transcripts by the enzyme RNA guanylyltransferase and 5’-phosphatase capping enzyme (RNGTT) or mRNA capping enzyme (CE)[17]. This capping process takes place in the nucleus, where RNGTT, along with RNA guanine methyltransferase (RNMT) and RNA guanine 7-methyl transferase activating subunit (RAMAC), adds the cap to the 5’ end of nascent RNA transcripts[18]. A population of cytosolic capping enzymes (cCE) has also been discovered. The cCE forms a complex with the cytoplasmic pool of RNMT, RAMAC, an unknown 5’RNA kinase, and the adaptor protein Nck1, providing a platform for the assembly of these components to make active cCE complex[19]. To study the function of cytoplasmic capping without interfering with the nuclear function, a stable cell line was generated expressing a dominant negative mutant of cCE where the catalytic lysine residue, was replaced with an alanine residue (K294A), to inhibit cytoplasmic recapping[20], [21]. cCE is responsible for adding a protective cap to specific mRNAs and lncRNAs, preventing uncapped RNAs from decaying. Thus, cytoplasmic capping is believed to maintain ‘cap homeostasis’ and regulate gene expression by influencing the stability of certain mRNAs and lncRNAs[21], [22]. The recapping process plays various roles in several biological processes like regulating drosophila development by interfering with hedgehog signaling [23], protecting the cap status of target mRNAs during arsenite-induced oxidative stress in osteosarcoma U2OS[24], and providing stability to the transcripts that control the mitotic cell cycle[21]. However, the post-transcriptional control of lncRNAs at the global transcriptomic level by the cCE is still elusive.

To induce hypoxia chemically in the cellular model, CoCl_2_ is widely used as a hypoxia mimetic reagent where expression of HIF1α in cell extract is used as an indicator of successful hypoxia induction[25]. In our previous study, we successfully established this CoCl2 induced hypoxic model and observed elevated expression of CE levels under CoCl_2_-induced hypoxic conditions[26]. This could potentially change the cellular response to hypoxia by stabilizing cCE-targeted lncRNAs. Interestingly, a significant upregulation of H19-009 expression was found in CoCl_2_-induced hypoxic cells, which was reduced in cells expressing K294A[26]. This reduction may be attributed to losing its 5’ cap structure during hypoxic stress. Without active cCE, it may not be recapped and could have been targeted by exoribonucleases. So, based on these results, we were interested in performing global RNA sequencing to identify the transcriptome-wide cCE-targeted, hypoxia-responsive novel lncRNAs in U2OS osteosarcoma cells. Additionally, we assessed the steady-state levels of selected lncRNAs in CoCl_2_ mimetic hypoxic conditions and CoCl_2_ untreated cells as well as in cells expressing or not expressing K294A. Our findings reveal that several of these lncRNAs are dysregulated under CoCl_2_-induced hypoxia and that cCE plays a crucial role in regulating their stability.

## Results & Discussion

### Differential Expression of lncRNAs in CoCl_2_-Induced hypoxic U2OS cells

In human solid tumors, hypoxia contributes to tumor growth, progression, and metastasis. To evaluate the effects of CoCl_2_-induced hypoxia on the expression profile of lncRNAs in the osteosarcoma cell line U2OS, we performed RNA sequencing. The following analysis identified 398 lncRNAs with significant differential expression under hypoxic conditions, comprising 186 upregulated and 212 downregulated lncRNAs (Figs 1A-B). The upregulation of numerous lncRNAs is closely linked to malignancy and angiogenesis[27], [28]. Therefore, we investigated the involvement of these dysregulated lncRNAs in various biological functions and pathways. Interestingly, the dysregulated lncRNAs are associated with different types of cancer as revealed by Gene Ontology (GO) enrichment analysis revealed that the upregulated lncRNAs in CoCl_2_-induced hypoxic U2OS cells were significantly associated with functional terms related to several cancers, including gastric, osteosarcoma, breast, colorectal, glioblastoma, and hepatocellular cancer (Fig 1C). Conversely, the downregulated lncRNAs were linked to functional terms such as astrocytoma, breast cancer, osteosarcoma, lymphoma, and melanoma (Fig 1D). Further analysis through the Kyoto Encyclopedia of Genes and Genomes (KEGG) pathway analysis identified 77 enriched pathways for upregulated lncRNAs. These pathways include the cell cycle, prostate cancer, MAPK signaling, cellular senescence, mTOR signaling, TNF signaling, and HIF-1 signaling pathways. The top 40 pathways, with significant p-values associated with the upregulated lncRNAs, are detailed in the accompanying Fig 1E. Similarly, 73 pathways were enriched for the downregulated lncRNAs, encompassing pathways such as the cell cycle, cellular senescence, mTOR signaling, chronic myeloid leukemia, and the HIF-1 signaling pathway (Fig 1F). The GO enrichment and KEGG pathway analyses show that the dysregulated lncRNAs associated with CoCl_2_-induced hypoxia are involved in various cancers and critical biological pathways.

**Figure 1:**
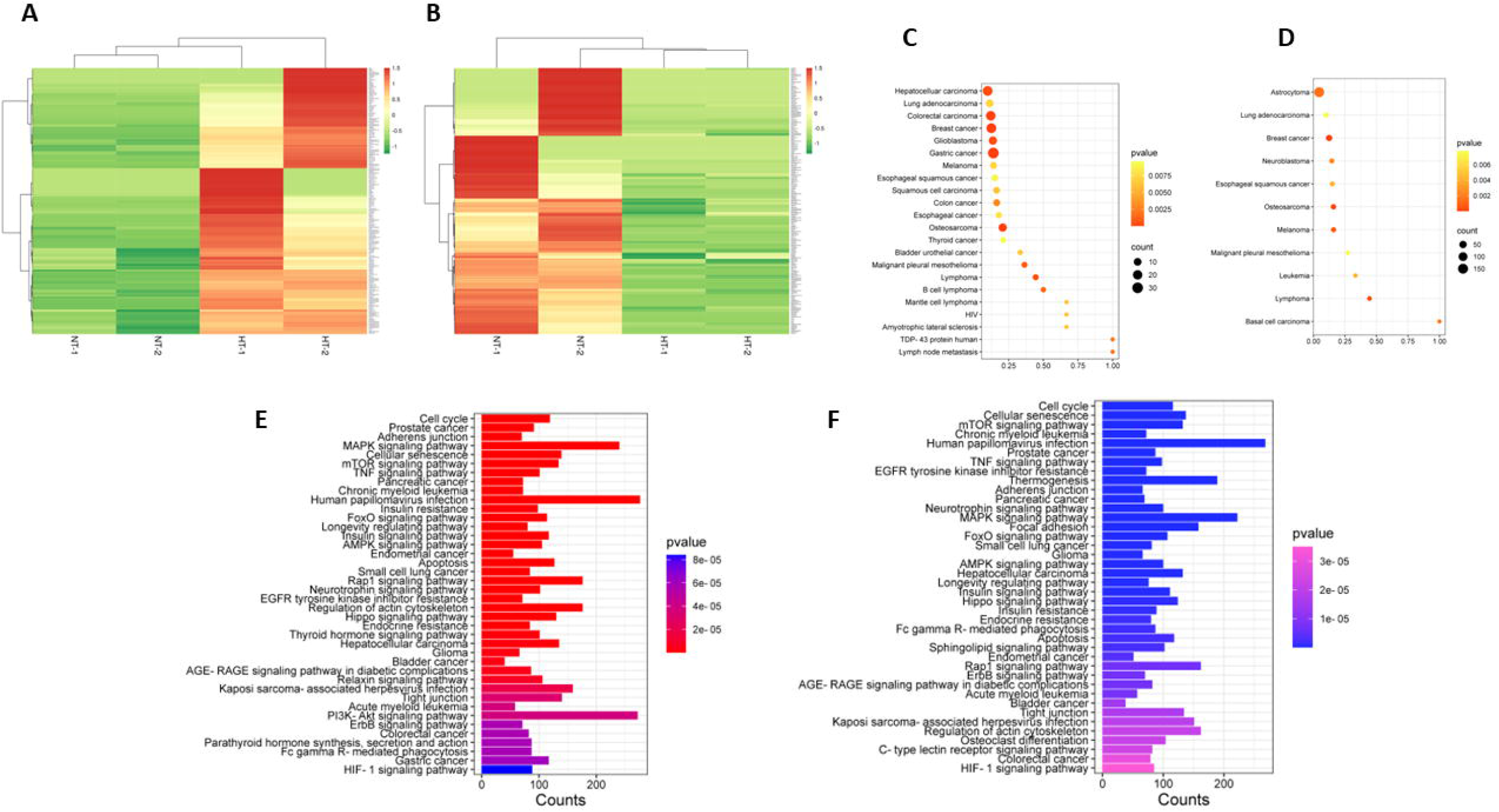
Expression profile and functional prediction of lncRNAs through RNA-sequencing under CoCl_2_-induced hypoxic conditions in U2OS cells: Clustered heatmap illustrating the upregulated **(A)** and downregulated **(B)** lncRNAs under hypoxic conditions. The expression level of each transcript (log2) is depicted according to the colour scale. The gradient scale indicates regions of upregulation (red) and downregulation (green) observed in this study. Rows denote lncRNAs, while columns represent the two sets of normoxic (NT1, NT2) and CoCl_2_ induced hypoxic (HT1, HT2) conditions. GO analysis revealed gene enrichment in different cellular biological processes with a 95% similarity index for upregulated **(C)** and downregulated **(D)** lncRNAs. KEGG analysis showing the top 40 significant signaling pathways associated with upregulated **(E)** and downregulated **(F)** lncRNAs.

### Experimental validation of RNA seq data

We have selected three upregulated lncRNAs, NEAT1-209, LUCAT1-211, and MSC-AS1-212, for further analysis due to their enrichment in GO clusters and associations with various diseases and pathways (Fig. 1). NEAT1 is a key component of paraspeckle nuclear bodies, influencing cell cycle regulation, proliferation, apoptosis, and tumor cell migration[29], [30], [31]. LUCAT1 is implicated in breast, ovarian, thyroid, and renal cancers, affecting tumor proliferation, invasion, and migration through diverse mechanisms[32], [33], [34], [35]. MSC-AS1 interacts with miR-425-5p, miR-325, and miR-373-3p to modulate ovarian cancer inhibition, colorectal cancer progression, and glioma cell growth and chemoresistance, respectively[36], [37], [38]. Since cytoplasmic RNA from U2OS cells was used for RNA seq experiments, we used the same fraction for validation. U2OS cells treated with or without CoCl2 were separated into nuclear and cytoplasmic components and the efficiency of the fractionation was examined by the presence of Lamin A/C in the nuclear fractions and GAPDH in the cytoplasmic fractions (Fig. 2A) To corroborate the findings from our RNA sequencing analysis, we isolated cytoplasmic RNA and performed qRT-PCR to assess the expression levels of selected lncRNAs that exhibited differential expression between ±CoCl_2_ added U2OS cells. Our results confirmed a significant upregulation in the steady-state expression of LUCAT1-211 and MSC-AS1-212 under hypoxic conditions (Fig. 2B) but not for NEAT1. This observation is consistent with the RNA sequencing data and suggests that CoCl2-induced hypoxia leads to the enhanced expression of specific lncRNAs in osteosarcoma cells. The potential explanation for the downregulation of NEAT1 is intriguing. NEAT1 exists in two isoforms: a shorter version approximately 3 kb in length (NEAT1_1) and a longer version around 23 kb (NEAT1_2).

**Figure 2:**
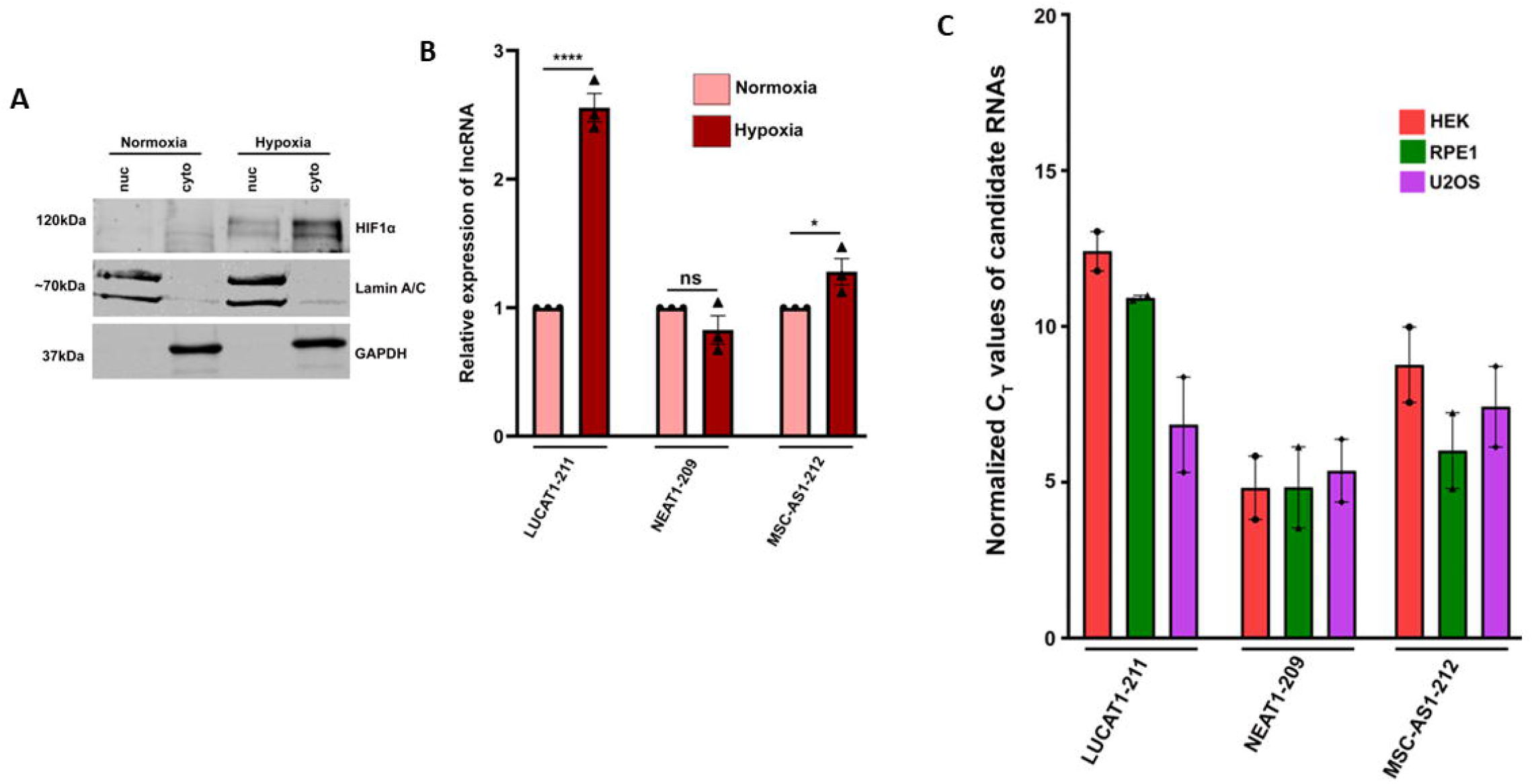
Validation of RNA Sequencing Under CoCl2-Induced Hypoxic Conditions: (A) Western blot showing nuclear and cytoplasmic fractions of U2OS cells. Presence of Lamin A/C in the nuclear fraction and GAPDH in the cytoplasmic fraction shows the purity of the fractions. Western blot analysis of the HIF1α (hypoxic marker) confirmed the effective induction of hypoxic conditions following a 24-hour treatment with CoCl_2_. **(B)** RT-qPCR analysis of cytoplasmic RNAs isolated from ± CoCl_2_ treated cells using gene-specific primers showed upregulated expression of LUCAT1-211 and MSC-AS1-212 in CoCl_2_ induced hypoxia, while the expression of NEAT1-209 was decreased. Statistical analysis was conducted using a two-tailed Student’s t-test. **(C)** RT-qPCR analysis showing variable expressions of LUCAT-211, MSC-AS1 and NEAT1-209 in cytoplasmic RNAs isolated from RPE1, U2OS and HEK293 cells. Values are expressed as mean ± standard deviation (SD) from three biological replicates. Statistical significance is denoted as follows: ns indicates non-significant; *P < 0.05; **P < 0.005; ***P < 0.0005; ****P < 0.0001, with n ≥ 3.

The longer isoform is crucial for paraspeckle formation and its levels increase in the absence of the arsenic resistance protein 2 (ARS2)[39]. Notably, the cytoplasmic isoform of ARS2 becomes more active only under arsenic stress conditions[40]. Given that we isolated the cytoplasmic RNA without arsenic stress, ARS2 should be absent under these conditions. Consequently, the longer NEAT1 isoform (NEAT1_2) would be expected to be elevated. However, our sequencing results have identified the shorter NEAT1 isoform. Moreover, pan-cancer analysis revealed that the expression of NEAT1 is downregulated in many cancers like breast invasive cancer, esophageal carcinomas, and pheochromocytoma & paraganglioma[41]. In the future, we could provide a more precise explanation.

These findings imply that the hypoxia signalling pathway may play a crucial role in regulating hypoxia-responsive lncRNAs, highlighting a potential mechanism through which hypoxia can influence gene expression in osteosarcoma. Subsequently, we investigated whether the selected lncRNAs are specific to osteosarcoma or if their expression varies according to cell lineage specificity. To this end, we utilized three distinct cell lines: the human retinal pigment epithelial line (RPE-1), which is a non-cancerous diploid cell line; the Human Embryonic Kidney (HEK) 293T, a transformed cell line; and the human osteosarcoma U2OS, a cancerous cell line. We isolated total RNA from these cell lines and employed qRT-PCR to analyze the expression levels of the selected lncRNAs. Our data show that the expressions of candidate lncRNAs are not restricted to cancerous cells (Fig. 2C). The variation in the expression of these lncRNAs across the cell line suggests different regulatory functions of these transcripts in different cells.

### HIF-1_α_ mediates the elevated expression of the selected lncRNAs

HIF1α is a well-established transcription factor activated under hypoxic conditions. In our CoCl_2_-induced hypoxia model, we observed an elevation in HIF1α protein levels (Fig. 3B). HIF1α binds to hypoxia response elements (HREs) with the consensus sequence R/CGTG located in the promoters or enhancers of its target genes. This binding facilitates the recruitment of coactivators such as p300 and CREB-binding protein, thereby initiating the transcription of various hypoxia-responsive genes. Given this background, we sought to determine whether the upregulation of these specific lncRNAs was mediated by HIF1α, the transcription factor induced by hypoxia. Utilizing the JASPAR database, we investigated the presence of the potential HIF1α binding sites within the selected lncRNAs. Our analysis revealed that LUCAT1 and NEAT1 contain 6 HIF1α binding sites each, while MSC-AS1 possesses two such sites (Fig.3A). To further elucidate the role of HIF1α, we employed PX-478, a specific inhibitor of HIF1α, to assess its effects. In the absence of CoCl_2_, HIF1α did not express, as anticipated. However, in CoCl_2_-induced hypoxic conditions without PX-478 treatment, HIF1α expression was detectable. Conversely, in hypoxic conditions treated with PX-478, no HIF1α bands were observed (Fig. 3B). Subsequently, we examined the expression of the selected lncRNAs under conditions where HIF1α was inhibited, using carbonic anhydrase 9 (CA9) as a positive marker for hypoxia[42] which showed upregulation in CoCl_2_ treated cells and downregulation in PX478 treated cells (Fig. 3C). In samples where HIF1α was inhibited, there was a noticeable reduction in the expression of LUCAT1 and MSC-AS1 compared to PX478 untreated, CoCl_2_-added cells (Fig. 3C). Following our previous data, NEAT1 did not show upregulated expression in hypoxic conditions and its expression level remained almost similar in HIF1α inhibited cells despite having 6 potential 6 HIF1α binding sites. Our results indicate that the increased expression of LUCAT1 and MSC-AS1 under CoCl_2_-induced hypoxic conditions is likely attributed to the direct binding of the HIF1α transcription factor to the HRE sites. This binding is effectively disrupted when HIF1α activity is blocked, underscoring the critical role of HIF1α in mediating the hypoxic response of these lncRNAs. Future studies should examine if LUCAT1 and MSC-AS1 bind directly to HIF1α in response to hypoxia and whether HIF1 binding sites of NEAT1 remained inactive.

**Figure 3:**
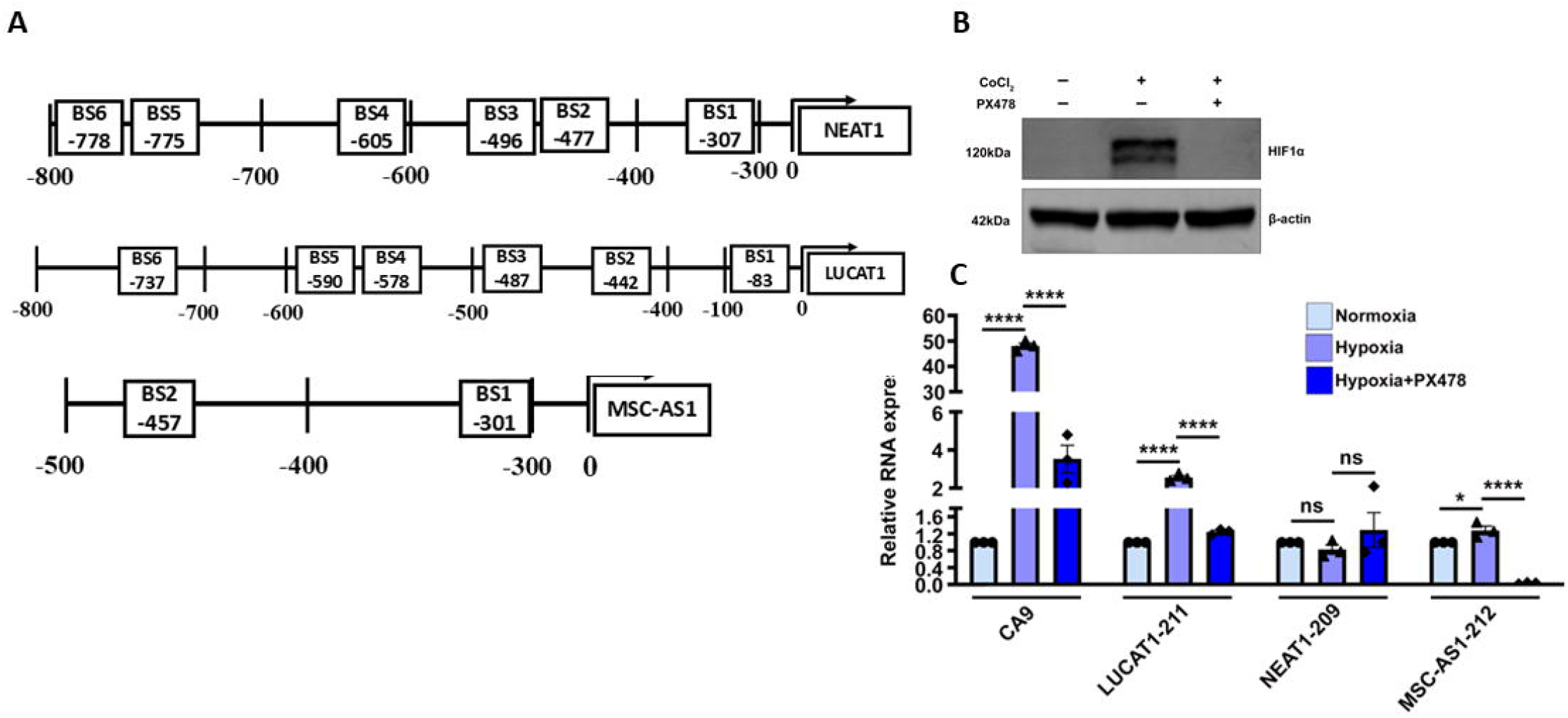
HIF1_α_ induces expressions of selected lncRNAs: **(A)** The selected lncRNAs possess multiple HIF1α binding motifs (R/CGTG) within their promoter regions, as illustrated in the schematic. **(B)** Western blot shows the addition of CoCl_2_ led to elevated expression of HIFα which was reduced after PX-478 treatment. **(C)** The RT–qPCR analysis of cytoplasmic RNA from the above treatment groups showed that treating U2OS cells with the HIF1α-specific inhibitor PX-478 under hypoxic conditions led to a significant decrease in the expression of LUCAT1-211 and MSC-AS1-212. CA9 serves as a known hypoxia-induced gene regulated by HIF1α[47]. Values are expressed as mean ± standard deviation (SD) from three biological replicates. Statistical significance is denoted as follows: ns indicates non-significant; *P < 0.05; **P < 0.005; ***P < 0.0005; ****P < 0.0001, with n ≥ 3.

### The hypoxia-responsive lncRNAs are regulated by mRNA cytoplasmic capping enzyme

Our previous study has demonstrated that specific lncRNAs can be targeted post-transcriptionally by the cCE[22]. Additionally, we have observed that LUCAT1 and MCS-AS1 are upregulated during CoCl_2_-induced hypoxia. We wondered if these lncRNAs selected for this study were regulated by cCE or not. To examine this, we used stable cells expressing ± K294A treated with or without CoCl_2_ for 24 hr followed by nuclear and cytoplasmic fractionation. Proteins and RNA were extracted from these cytoplasmic fractions. To verify the expression of the induced proteins, we performed western blot analysis utilizing cytoplasmic proteins extracted from cells stably expressing either the vector control or the K294A construct. The efficiency of fractionation was measured by the presence of Lamin A/C in the nuclear fraction and GAPDH in the cytoplasmic fraction (Fig. 4A). Our data showed expression of Myc-tagged K294A protein only in the cytoplasmic fractions from U2OS cell line expressing K294A (Fig. 4A). Subsequently, we assessed the steady-state expression levels of candidate lncRNAs in cytoplasmic fractions isolated from cells expressing vector (Control) and K294A (K294A) using RT-qPCR. The expression levels of LUCAT1, NEAT1, and MSC-AS1 were significantly diminished in K294A cells compared to Control cells (Fig. 4B). This reduction in the expression of selected HRLs in K294A cells may be attributed to the inactivity of the cytoplasmic capping enzyme. Due to the absence of active cCE, it is plausible that the selected HRLs present in the cytoplasm could not undergo recapping, rendering them vulnerable to exoribonuclease-mediated degradation, thereby resulting in their decreased expression in K294A cells.

**Figure 4:**
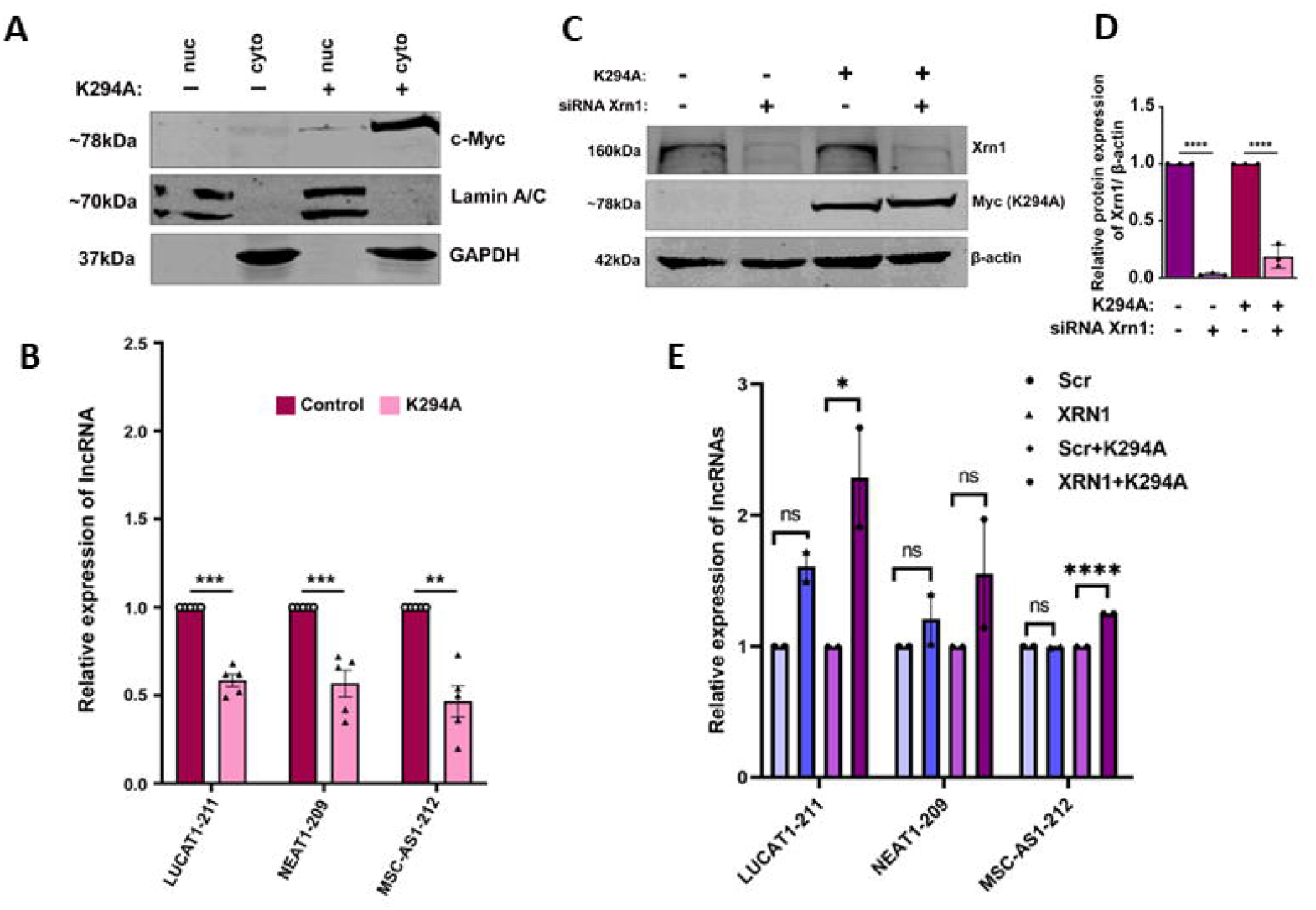
The hypoxia-responsive lncRNAs are regulated by mRNA cytoplasmic capping enzyme: **(A)** Western blot analysis of U2OS cells expressing ± K294A showed Myc-tagged K294A expression. The quality of nuclear and cytoplasmic fractionation was validated by the detection of nuclear marker Lamin A/C and cytoplasmic marker GAPDH. **(B)**The steady-state expression of NEAT1-209, LUCAT1-211, and MSC-AS1-212 was significantly decreased in K294A-expressing cells. Statistical analysis was conducted using a two-tailed Student’s t-test. **(C)** The impact of siRNA-mediated Xrn1 knockdown in U2OS cells on the stability of the Xrn1 exoribonuclease was evaluated using Western blot analysis. Simultaneously, the Western blot for Myc verified the expression of the K294A variant. β-Actin served as the internal control. **(D)** The relative expression levels of Xrn1 protein, following transfection with Xrn1 siRNA, were quantified by normalization against β-Actin protein expression in ±K294A cells. **(E)** The steady-state levels of NEAT1-209, LUCAT1-211, and MSC-AS1-212 were significantly elevated following the transfection of siRNA Xrn1 into K294A U2OS cells. Statistical analysis was conducted using the two-way ANOVA. Values are expressed as mean ± standard deviation (SD) from three biological replicates. Statistical significance is denoted as follows: ns indicates non-significant; *P < 0.05; **P < 0.005; ***P < 0.0005; ****P < 0.0001, with n ≥ 3.

Therefore, we hypothesized that the inhibition of the cCE would result in a decrease in the steady-state levels of cCE-targeted lncRNAs. This hypothesis is based on the premise that uncapped monophosphorylated RNAs, lacking the recapping function of cCE, would become substrates for degradation by the Xrn1 exonuclease. Notably, Xrn1 is a 5’-3’ exoribonuclease that plays a pivotal role in RNA metabolism, crucial for maintaining cellular homeostasis and preventing the accumulation of aberrant RNA species. To investigate the impact of Xrn1 on the stability of cCE-targeted lncRNAs, we downregulated expression of Xrn1 using siRNA specific to Xrn1 in Control and K294A cells (Fig. 4C). Western blot analysis and relative protein quantification showed more than 80% reduction in Xrn1 levels in both control and K294A cell lines when treated with Xrn1 siRNA compared to the scrambled control siRNA. Myc-K294A was expressed only in K294A cells, but not in control cells (Fig. 4C - 4D).

Next, to determine the impact of Xrn1 knockdown in the steady state levels of the cCE-targeted lncRNAs, cytoplasmic RNA was extracted from these cells and RT-qPCR was performed. Our findings revealed an upregulation of LUCAT1-211 and MSC-AS1-212 in K294A cells when Xrn1 was knocked down (Fig. 4E). Similar trend was also noticed for NEAT1 but the data remained insignificant (Fig. 4E). These results suggest that the steady-state levels of these lncRNAs are negatively regulated by Xrn1 exonuclease in the absence of the cytoplasmic capping enzyme. This observation implies that cCE may play a role in post-transcriptionally regulating these lncRNAs, thereby contributing to their stability.

## Conclusion

This study identified 306 lncRNAs associated with CoCl_2_-induced hypoxic conditions in U2OS osteosarcoma cells. Our findings also suggest that cCE may regulate a few HRLs at the post-transcriptional level. Future studies should examine more HRLs as obtained from the RNA seq data and probable regulation of cCE in response to chemical or physically induced hypoxia. Another possibility could be the regulation of hypoxic signalling pathways by cCE which in turn influence the TME and could be studied in the future. Overall, the findings from this study suggest that cCE could emerge as a pivotal factor in modifying cellular responses to hypoxia by stabilizing cCE-targeted lncRNAs.

## Materials and methods

### Cell culture and hypoxia induction by CoCl_2_

The U2OS human osteosarcoma cell line was acquired from the American Type Culture Collection (ATCC). It was cultured in 1X DMEM (Gibco, cat. no. 11965092) with 10% fetal bovine serum (Himedia, cat. no. RM1112) and 1% penicillin-streptomycin (Gibco, cat. no. 15140-122) under 5% CO2 at 37°C in a humidified Eppendorf incubator. Mycoplasma contamination was assessed using a PCR detection kit (Sigma-Aldrich, cat. no. MP0035) to ensure a contamination-free environment. To induce hypoxia *in vitro*, cells were treated with 200 µM Cobalt chloride (Sigma Aldrich, cat no# C8661) dissolved in DPBS (Gibco, cat no# 14040182). After 24 hours, cells were harvested for RNA/protein extraction. The HIF1α inhibitor PX478 (MedChemExpress, cat no #HY-10231) was prepared as a 10mM stock in nuclease-free water. Cells were then treated with 40µM PX478 for hours to inhibit HIF1α activity.

### Generation of stable cell lines

In this study, U2OS cells were initially transfected with the pLenti6/TR vector (obtained from Dr. Shubhra Majumder, Presidency University). For the selection of clonal cells that stably express the Tetracycline repressor (Tet-R), Blasticidin S Hydrochloride (Sigma Aldrich, cat no#15205) was employed. Multiple clones were successfully isolated and subsequently analyzed for the Tet-R protein’s expression. This analysis was conducted utilizing the western blotting technique, employing a specific antibody targeting Tet-R (Novus Biologicals cat no# NB600-234). Among the clones, the one expressing the highest level of the repressor was subsequently transfected with the pcDNA4-TO-bio-myc K294A construct, as described by Mukherjee et al. (2012)[21]. To establish stable cell lines, Zeocin (Invitrogen, cat no# R25001) was utilized for selection. Following this, Doxycycline was administered at a concentration of 1 µg/ml (Sigma Aldrich, cat no# D5207) to induce the expression of the K294A protein. The cell line that exhibited the highest expression of the K294A protein was then selected for further experimental analyses. As control cells, U2OS cells stably expressing the pLenti6/TR were used throughout the study.

### Xrn1 knockdown

U2OS Tet-R and K294A cells were transfected with Pre-validated Silencer Select siRNA targeting the coding region of XRN1 (Ambion, assay ID: s29016) at a final concentration of 10 nM, using Lipofectamine RNAiMAX (cat no# 13778-075; Invitrogen) as per manufacturer’s instruction. 24 hours post-transfection, the transfection mix containing media was replaced with a fresh complete medium. After 48 hours of post-transfection, K294A cells were treated with doxycycline (1 μg/ml) to induce K294A expression. 24 hours post doxycycline treatment cells were harvested for RNA and protein extraction. For control of knockdown experiments, cells expressing ±K294A cells were transfected with scramble siRNA and incubated for same time. The XRN1 knockdown efficiency was measured by western blotting with rabbit anti-Xrn1 antibody (Thermo scientific cat no# PA5-57110).

### Nuclear & Cytoplasmic fractionation

U2OS cells were collected using ice-cold DPBS, and the resulting cell pellet was obtained by centrifugation at 1000g for 5 minutes at 4°C. The pellet was subsequently resuspended in a cytoplasmic lysis buffer composed of 20 mM Tris-HCl (pH 7.5), 10 mM NaCl, 10 mM MgCl2, 10 mM KCl, and 0.2% NP-40, with the addition of 1 mM phenylmethylsulfonyl fluoride (PMSF). The buffer was further supplemented with RNaseOUT (Invitrogen, cat no# 10777019) and a protease inhibitor cocktail (Roche, cat no# 4693159001) to prevent RNA and protein degradation. The cell suspensions were maintained on ice for 10 minutes and gently agitated at intervals. Following this incubation, the suspensions were centrifuged at 1000g for 10 minutes at 4°C. The supernatant, containing the cytoplasmic fraction, was carefully transferred to a new tube for subsequent analysis or processing, and the pellet containing nuclei. The nuclear lysate was prepared by dissolving the nuclei pellet collected in nuclear lysis buffer (25 mM Tris-HCl pH 7.5, 150 mM NaCl, 1% triton X 100, 0.5% sodium deoxycholate, 1% SDS) supplemented with RNaseOUT and PMSF. The suspension was kept in ice for 30 minutes with intermittent vortex, then centrifuged at 10,000g for 10 minutes at 4°C. The supernatant containing the nuclear fraction was carefully transferred to a fresh tube for further analysis and the pellet was discarded.

### Cytoplasmic RNA extraction

Cytoplasmic RNA was extracted from the cells utilizing the TRIzol reagent (Invitrogen, cat no# 15596026), adhering strictly to the manufacturer’s protocol. Subsequently, the extracted RNAs underwent treatment with DNase I (Thermo Scientific, cat no# EN0521) to eliminate any contaminating DNA. Following this, a purification process was conducted using a mixture of phenol, chloroform, and isoamyl alcohol (Sisco Research Laboratories Pvt Ltd, cat no# 69031) to ensure the integrity and purity of the RNA samples.

### Next Generation Sequencing

The isolated cytoplasmic RNA from cells treated with or without CoCl_2_ have been sent for Next Gen sequencing using Illumina platform where the standard protocol has been followed for library preparation. Extracted RNA quantity is checked on Qubit flurometer (Thermofisher, cat no#Q33238) using RNA HS assay kit (Thermofisher, cat no #Q32851) following the manufacturer’s protocol. The library preparation was carried out using TruSeq® Stranded total RNA kit (Illumina, cat no#15032611). Final libraries were quantified using Qubit 4.0 fluorometer (Thermofisher #Q33238) using DNA HS assay kit (Thermofisher, cat no #Q32851) following manufacturer’s protocol. All libraries can be further processed for the Sequencing using Illumina platform. Long non-coding RNA transcript sequences for *Homo Sapiens* (Source: GENCODE; Release 42 (GRCh38.p13) was obtained from GENCODE (https://www.gencodegenes.org/human/). The transcript-to-transcript mapping file was prepared by extracting the FASTA header of the long non-coding transcript FASTA file. Long non-coding RNA transcript file was indexed using Kallisto v0.48[43]. Processed samples were quantified against the indexed human Long non-coding RNA transcript using quant utility of Kallisto with default parameters. Normalization and differential expression analysis were performed using DESeq2[44], The data will be deposited soon in the GEO database.

### Quantitative real-time PCR analysis

1 µg of cytoplasmic RNA were reverse transcribed utilizing the iScript cDNA Synthesis Kit (Bio-Rad, cat no# 1708890), following the manufacturer’s guidelines. Subsequently, a 1:1 dilution of the resulting cDNA was prepared for real-time PCR analysis. This analysis employed the SsoAdvanced Universal SYBR Green Supermix (Bio-Rad, cat no# 172-5271) on a CFX Connect Real-Time PCR Detection System (Bio-Rad). The PCR amplification was conducted under the following thermal cycling conditions: an initial denaturation step at 95°C for 30 seconds, succeeded by 40 cycles consisting of denaturation at 95°C for 10 seconds and annealing/extension at 60°C for 15 seconds. Following the amplification, a melting curve analysis was performed by incrementally increasing the temperature from 65°C to 95°C in 0.5°C steps, with each step lasting 2 seconds, to verify the specificity of the PCR products. The housekeeping gene RPLP0 served as the internal control to normalize the data. The sequence of the gene specific primers are listed in Supplementary Table-1. The relative expression levels of the target genes were quantified using the comparative CT method, also known as the ΔΔCt method. All experiments were conducted with a minimum of three independent biological replicates to ensure the reliability and reproducibility of the results.

### Western blotting analysis

Cells were lysed using a protein lysis buffer composed of 50 mM Tris-Cl at pH 8.0, 150 mM NaCl, and 1% NP-40, with the addition of 1 mM PMSF and a protease inhibitor cocktail (Roche, cat no# 4693159001) to prevent protein degradation. Protein quantification was performed utilizing the Rapid Gold BCA Protein Assay Kit (Pierce, cat no# A53226). Protein extracts, ranging from 30 to 60 µg, were separated by 8-10% SDS-PAGE under reducing conditions and subsequently transferred onto Immobilon-FL PVDF membranes (Millipore, cat no# IPFL00010). The membranes were blocked with 3% skimmed milk in phosphate-buffered saline (PBS) for 30 minutes at room temperature. Following blocking, the membranes were incubated overnight at 4°C with constant shaking in the respective primary antibody dilutions. The primary antibodies used included: 1:1000 rabbit anti-HIF1α (Cell Signaling Technology, cat no# 36169), 1:1000 mouse anti-β actin (Santa Cruz Biotechnology, cat no# SC-517582), 1:1000 rabbit anti-CE (Novus Biologicals, cat no# NPB1-49973), 1:2000 mouse anti-GAPDH (Novus Biologicals, cat no# 2D4A7), 1:2000 rabbit anti-CA9 (Cloud-clone Corp, cat no# PAD076Hu01), 1:2000 mouse anti-lamin (DHSB, cat no# MANLAC1(4A7)), and anti-Xrn1 (cat no# PA5-57110).

Post-primary antibody incubation, the membranes underwent three washes with PBS-T. Secondary antibodies were then applied, including donkey anti-Mouse DyLight 800 (Cell Signaling Technology, cat no# 5257) and donkey anti-Rabbit 680 (Invitrogen, cat no# A10043), both diluted at 1:10000 in PBS-T. The blots were scanned using the Odyssey CLx Imaging System (Li-Cor Inc.), and band intensities were quantified using ImageJ software (NIH, USA).

### Different expression analysis of lncRNAs and heatmap generation

The differential expression analysis of the lncRNAs was conducted using the DESeq2 software. lncRNAs were identified as differentially expressed if they met the criteria of having a minimal false discovery rate (FDR) and an absolute fold change of greater than 1. Subsequently, these differentially expressed lncRNAs underwent clustering through the ClustVis R package (https://github.com/taunometsalu/ClustVis), which facilitated both Principal Component Analysis and the generation of heatmaps[45].

### Gene Ontology analysis and KEGG enrichment

GO enrichment analysis identifies GO terms that are significantly enriched in differentially expressed genes when compared to the genomic background. For the analysis of the lncRNAs, the TLSEA tool was utilized with a similarity coefficient of 0.95. This tool employs graph representation learning methods to extract low-dimensional vectors of lncRNAs within two functional annotation networks, facilitating a deeper understanding of lncRNA functions (http://www.lirmed.com:5003/tlsea).

Genes often interact with one another to fulfil specific biological roles. To gain further insights into the biological functions of genes, KEGG pathway-based analysis is employed.

This approach provides a comprehensive understanding of the pathways in which genes are involved. The visualization and enrichment analysis were conducted using the NcPath database[46], which offers a robust platform for such analyses (https://github.com/Marscolono/NcPath/tree/main). A p-value of less than 0.05 was considered indicative of statistically significant enrichment, ensuring the reliability and validity of the findings.

### Statistical analysis

The statistical analysis was calculated by performing Two-tail Student’s t-tests for two groups and one-way ANOVA for multiple groups using GraphPad Prism 9 Software (GraphPad Software, Inc). All the data were represented as ±SD of at least three independent biological replicates (n=3). Data with p<0.05 were considered statistically significant.

## Acknowledgments

This work was supported by extramural grants CRG/2019/006427 and CRG/2023/006384 from the Department of Science and Technology, Government of India to Dr. Chandrama Mukherjee. The authors also thank UGC, Government of India for a UGC-SRF fellowship to Mr. Safirul Islam and the Department of Biotechnology, India for a Ramlingaswami Re-entry fellowship to Dr. Chandrama Mukherjee. The authors thank the Institute of Health Sciences, Presidency University for the departmental instrument facility, and Presidency University for necessary infrastructural support.

