## Supplementary Table 1 for "Identification of transcriptome-wide cobalt chloride-induced hypoxia-responsive long noncoding RNAs regulated by cytoplasmic mRNA capping enzyme"

| **Primer Name** |  | **Sequence (5'-3')** |
| --- | --- | --- |
| NEAT1-209 | Forward | GCTCTTGCATAGCTGAGCGA |
|  | Reverse | TGATCATTTCCAGGGCTGCTG |
| MSC-AS1-212 | Forward | CAGACTGAGAGCCCAATG |
|  | Reverse | GGAACTTTTACTCGTGGC |
| LUCAT1-211 | Forward | GAGTAGCTGGGACTACAGGC |
|  | Reverse | ACAGGCACGCTAAGTCTCATC |
| RPLPO | Forward | GGAGAAACTGCTGCCTCATATC |
|  | Reverse | CAGCAGCTGGCACCTTATT |
| Supplementary Table-1: Sequences of primers used in this study | | |
